# Microbial community assembly differs by mineral type in the rhizosphere

**DOI:** 10.1101/128850

**Authors:** Thea Whitman, Rachel Neurath, Adele Perera, Daliang Ning, Jizhong Zhou, Peter Nico, Jennifer Pett-Ridge, Mary Firestone

## Abstract

Inputs of root carbon (C) fuel growth of nearby soil microorganisms. If these microbes associate with soil minerals, then mineral-microbiome complexes near roots could be a gateway towards stabilization of soil carbon and may influence the quantity and quality of persistent SOM. To investigate the interactions between roots, soil minerals, and microbes, we incubated three types of minerals (ferrihydrite, kaolinite, quartz) and a native soil mineral fraction near roots of a common Californian annual grass, *Avena barbata,* growing in its resident soil. We followed microbial colonization of these minerals for 2.5 months – the plant’s lifespan. Bacteria and fungi that colonized mineral surfaces during this experiment differed across mineral types and differed from those in the background soil, implying microbial colonization was the result of processes in addition to passive movement with water to mineral surfaces. Null model analysis revealed that dispersal limitation was a dominant factor structuring mineral-associated microbial communities for all mineral types. Once bacteria arrived at a mineral surface, capacity for rapid growth appeared important, as ribosomal copy number was significantly correlated with relative enrichment on minerals. *Glomeromycota* (a phylum associated with arbuscular mycorrhizal fungi) appeared to preferentially associate with ferrihydrite surfaces. The mechanisms enabling colonization of soil minerals may be foundational to the overall soil microbiome composition and partially responsible for the persistence of C entering soil via plant roots.

## Introduction

Plant roots and the microorganisms that surround them are critically important to soil C stabilization, as roots are the primary source of stabilized organic C in soil (Dijkstra and Cheng, 2007; Clemmensen *et al.*, 2013; Drake *et al.*, 2011; Treseder and Holden, 2013). This input of organic carbon fuels the growth of mineral-associated microbes that drive many critical soil processes, including mineral weathering (Banfield *et al.*, 1999; Uroz *et al.*, 2009), aggregate formation (Six and Paustian, 2014) and the cycling of mineral-sorbed organic matter (Saidy *et al.*, 2014; Schmidt *et al.*, 2011). Mineral sorption of soil organic matter (SOM) is thought to play an important role in limiting the microbial availability of soil C and contributing to the persistence of C in soils (Keiluweit *et al.*, 2015; Schmidt *et al.*, 2011), but is not well understood in the rhizosphere, where levels of organic inputs, exudates, and microbial activity are high (Kuzyakov and Blagodatskaya, 2015). Understanding the processes that control such mineral-SOM-microbe interactions is essential to understanding how microbial communities affect soil C cycling (Kallenbach *et al.*, 2016).

Given that one gram of soil may contain a billion bacterial cells, the patchiness of soil microbial communities can be surprising (Raynaud and Nunan, 2014). In most soils, mineral surfaces are not fully colonized by microbes (Nunan *et al.*; Ranjard and Richaume, 2001; Vos *et al.*, 2013), and are not “saturated” with organic matter (Kögel-Knabner *et al.*, 2008; Miltner *et al.*, 2011; Lehmann *et al.*, 2007). Fresh mineral surfaces are constantly regenerated in surface soils through the dynamic processes of mineral weathering and formation. While the canonical role of lichens in rock colonization and subsequent soil formation is well described (Chen *et al.*, 2000; Raab *et al.*, 2012; Cooper and Rudolph, 1953; Hodkinson *et al.*, 2002)), we know little about the first inhabitants of minerals as they form within the soil (Hutchens, 2009). In addition to the formation of new microhabitats through mineral weathering, frequent disturbances, ranging from large-scale climatic changes (Pold and DeAngelis, 2013) to the millimeter-scale incursion of roots (Belnap *et al.*, 2003), regularly “reset” microscale communities (DeAngelis *et al.*, 2008). These disruptions ensure that meaningful “stable states” or “climax communities” are rare, and microbial colonization processes are likely important determinants of soil microbial community composition. Studying microbial colonization of “fresh” soil minerals (*i.e.*, minerals free of SOM and cells) can provide insight into microbial community assembly in the “mineralosphere” (Uroz *et al.*, 2009) and in soils as a whole. In addition, microbial interactions with minerals have important implications for mineral dynamics, affecting metal speciation, toxicity, mobility, mineral formation, and mineral dissolution (Gadd, 2010).

Surface attachment confers important advantages for soil microorganisms, including protection from predation, access to nutrients or energy sources, and provision of a substrate for biofilm formation or other density-dependent phenomena (Hutchens, 2009; Uroz *et al.*, 2015). However, soil minerals can provide much more than simply an attachment surface. Different minerals offer specific chemical or physical environments – *e.g.*, varying in surface area, redox status, or chemical composition (Banfield and Hamers, 1997) – which may regulate the degree of microbial colonization and even community composition. For example, (Hutchens *et al.*, 2010) found significantly different bacterial and fungal communities colonized different granitic minerals within the same exposed rocky outcrop. However, very few mineral colonization studies have been done in a soil context (Uroz *et al.*, 2015). Uroz *et al*. (2012) found that different communities colonized minerals (pure apatite, pure plagioclase and a mix of phlogopite-quartz) after four years of burial in acidic forest soils, while Wilson *et al*. (2008) found that while some minerals were more intensively colonized, magnetically-separated Fe-/Mg- minerals *vs*. volcanic glass or K-feldspar minerals had similar microbial community composition in a volcanic soil. Clearly, the effect of soil mineralogy on microbial colonization of fresh minerals and the mechanisms that control their community assembly processes require additional investigation (Uroz *et al.*, 2015). In our study, we explored these phenomena within a rhizosphere context, where altered chemical and resource characteristics create a unique environment.

The factors that shape community assembly (Maherali and Klironomos, 2007; Keddy, 1992; Drake, 1990; Tilman, 2004) and community succession within new habitats (Hodkinson *et al.*, 2002; Gleason, 1939; Young *et al.*, 2001) have long been studied for macrobiota. Similar principles may be applied to understand assembly of microbial communities. For example, in a fluid ecosystem, Zhou *et al*. (2014) evaluated the concepts of deterministic versus stochastic processes, showing that both processes played important roles, but their relative importance varied over time. In our study of microbe-scale colonization of fresh mineral surfaces within the rhizosphere, we considered a number of central questions. Are community members passively transported or do they actively move to new microhabitats? Once microbes arrive at a new microhabitat, does selection favor certain organisms from the source community, or is a novel community drawn indiscriminately from the surrounding rhizosphere soil? We investigated which microbes colonize “fresh” mineral surfaces in the soil, what community assembly processes determine initial community composition in the mineralosphere, and the characteristics that make microbes strong colonizers of mineral surfaces in rhizosphere soil. To address these questions, we incubated fresh minerals commonly found in our study soil (quartz, kaolinite, and ferrihydrite) as well as density-fractionated native soil minerals (“heavy fraction”) in soil microcosms planted with the annual grass *Avena barbata* for up to 2.5 months. We then assessed how root presence and different minerals selected for distinct fungal, bacterial, and archaeal communities in the soil. We tested whether the microbial composition of the fresh minerals came to resemble that of the surrounding soil (communities dominated by homogenizing selection and homogenizing dispersal), or whether some degree of community structuring was driven by differences in minerals (variable selection or dispersal limitation). We tested whether the microbial composition of the fresh minerals came to resemble that of the surrounding soil (communities dominated by homogenizing selection and homogenizing dispersal), or whether some degree of community structuring was driven by differences in minerals (variable selection or dispersal limitation).

## Materials and Methods

### Experimental design

*Avena barbata* (wild oat) plants were grown in a soil from a California annual grassland that supports *A. barbata* as a dominant grass species. The soil is a fine-loamy, mixed, active, mesic Typic Haploxeralf (properties described in Supplementary Table 1) collected from 0-10 cm depth in a pasture at the UC Hopland Research and Extension Centre (38.992938° N, - 123.067714° W). It was sieved to < 2 mm and packed at field density (1.21 g cm^-3^) into mesocosms with a removable side panel (described in (DeAngelis *et al.*, 2008; Jaeger *et al.*, 1999). Plants (4 per microcosm - equivalent to field density) were grown under 14 hr full spectrum light, at 14%_vwc_ moisture, and 400 ppm CO_2_ for 1 month in the main chamber, after which the microcosms were opened, the side panel was removed, and 10 mineral bags (3 each of ferrihydrite, quartz, and kaolinite, and one of the heavy fraction; description follows) were placed directly on top of the roots and soil in a randomized order, covered with additional fresh soil, and the microcosms were resealed (Figure 1). Five microcosms were opened and mineral bags were destructively harvested after 1 month, 2 months, and 2.5 months of incubation, at which point plants were beginning to show signs of senescence. We also collected soil at this time. Soil was separated from coarse roots and passed through < 2 mm sieve to homogenize it, and then sub-sampled and preserved for analysis. All mineral and soil samples were immediately frozen on dry ice and placed in a -80 °C freezer for storage within the day.

**Figure 1.**
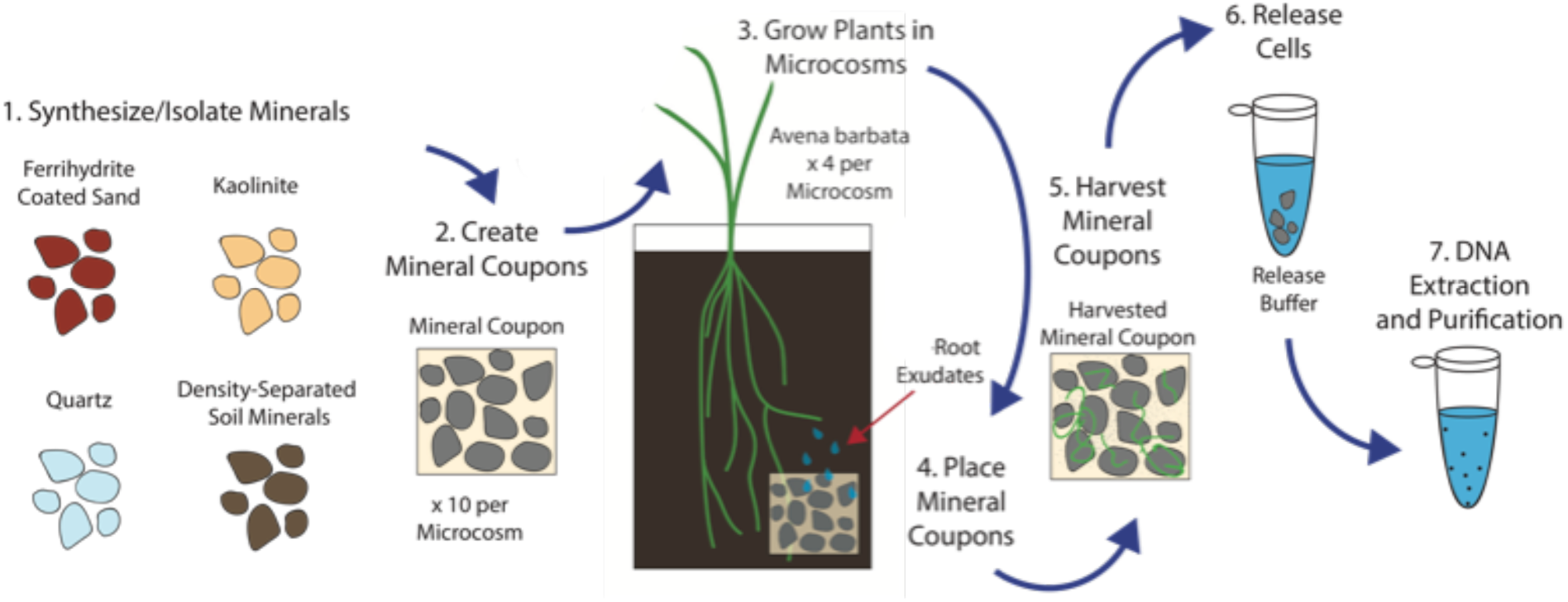
Experimental design conceptual figure. Ferrihydrite, kaolinite, and quartz minerals with pure, C-free surfaces and density-separated soil minerals were incubated in soil microcosms with growing *Avena barbata*. Minerals were harvested and immediately transferred to a cell release buffer and then DNA was extracted and purified for sequencing and qPCR.

Mineral preparation and properties are described in detail in Neurath *et al*. (*in prep*) and are summarized in Table 1. Briefly, X-ray diffraction (XRD) analysis of Hopland soil was used to identify the dominant clay mineral and iron oxide used in this study: kaolinite and ferrihydrite, respectively. Hopland soil also contains quartz, which we used as a “control” mineral due to its low surface area and less-reactive surface. Density fractionation (modified method by Sollins *et al.* (2006)) was used to separate the “heavy fraction” (>1.75 g-cm^-3^) component of Hopland soil from free light and occluded light fractions. The heavy fraction was then lyophilized before use in this experiment. Quartz sand was acid washed in 10% HCl. Ferrihydrite-coated quartz was synthesized in the lab, with Al and Si inclusion to better represent a natural ferrihydrite mineral. Kaolinite was mixed with quartz at a 1:1 per mass ratio to moderate potential clumping effects of pure clay. Minerals were sealed in 18 μm nylon mesh bags measuring 5 × 5 × 0.2 cm, with a single mineral type in each bag: density-separated heavy fraction; quartz; ferrihydrite-coated quartz (“ferrihydrite”); and the 50:50 kaolinite:quartz mix (“kaolinite”). The quartz, ferrihydrite, and kaolinite minerals had an initial C content of 0%, while the heavy fraction had an initial C content of 1.6%.

**Table 1.** Mineral properties

| Property | Quartz | Pure Kaolinite* | Ferrihydrite | Heavy fraction |
| --- | --- | --- | --- | --- |
| Chemical formula | $\text{SiO}_4$ | $\text{Al}_2\text{Si}_2\text{O}_5(\text{OH})_4$ | $\text{Fe}(\text{OH})_3$ | NA |
| Source of mineral | Purchased from Sigma-Aldrich, (274739) | Kaolinite purchased from the Clay Minerals Society (K-Ga2) | Synthesized in lab following (Hansel *et al.*, 2003) | Extracted from soil using modified protocol from (Sollins *et al.*, 2006) |
| BET surface area<sup>‡</sup> ($\text{m}^2 \text{ g}^{-1}$) | 0.01-0.05** | 20.48 | 4.8 | 2.68 |
| Particle size range | 297-210 $\mu\text{m}$ | Mostly $< 2\mu\text{m}$ | Coated quartz | Not determined |
| Initial C (%) | Negligible | Negligible | Negligible | 1.6 |
| Predicted relative charge density | Very low | Low | High | Intermediate |
| Primary or secondary mineral? | Primary | Secondary | Secondary | NA |

<sup>‡</sup> $\text{N}_2$ analysis gas
\* Kaolinite was used in a 50:50 mixture with quartz
\*\* Quartz surface area was too low to measure using the above techniques with $\text{N}_2$ gas.
Estimated from the literature ((Xu *et al.*, 2009; Mekonen *et al.*, 2013) and mesh size.

### DNA extraction

At harvest, minerals and bulk soil samples were transferred to Whirl-pak bags, frozen on dry ice, and stored at -80°C. Anticipating potential difficulty in DNA desorption from the ferrihydrite minerals in particular, we used a sterile cell release buffer of Tween20 (5 g L^-1^) and sodium pyrophosphate decahydrate (1 g L^-1^) (Supplementary Note 1). Released cells were processed using a modified phenol-choloroform DNA extraction protocol (Griffiths *et al.*, 2000; Shi *et al.*, 2015). Briefly, samples received 500 μL 5% CTAB / 0.7M NaCl / 240 mM K-PO_4_ at pH 8, 500 μL of 25:24:1 phenol/chloroform/isoamyl alcohol, and lysing matrix E beads (MP Biomedicals, Santa Ana, CA). Tubes were shaken on a FastPrep (MP Biomedicals, Santa Ana, CA) for 30 s on speed 5.5. After centrifuging at 4 °C, the aqueous phase was transferred to 2 mL phase-lock gel heavy tubes (5 Prime), where they received an equal amount of 24:1 chloroform/isoamyl alcohol, were mixed, centrifuged, and then the aqueous phase was transferred to 1 mL 40% PEG 6000 / 1.6 M NaCl, where DNA precipitated for 1 h. Extracted samples were then re-extracted with 500 μL CTAB mixture, with the resulting aqueous extract added to the PEG 6000 tubes, along with 1 μL linear acrylamide as a co-extractant. Samples were then centrifuged for 20 minutes at 4°C, supernatant removed, and then DNA pellets rinsed with 70% EtOH, air-dried, and resuspended in 50 μL RNAase-free water, and frozen at -80°C. DNA was quantified using a Quant-iT PicoGreen double stranded DNA assay kit (Invitrogen, Carlsbad, CA) and a BioRad iCycler (BioRad Laboratories, Hercules, CA).

### Quantitative PCR

The 16S rRNA gene and ITS DNA copies in each sample were determined using quantitative PCR (qPCR) with primer sets EUB338/EUB518 for bacteria and 5.8S/ITS 1f for fungi (Supplementary Table 2) (Fierer *et al.*, 2005), using a BioRad iCycler (BioRad Laboratories, Hercules, CA) and SSOFast EvaGreen Supermix (BioRad Laboratories, Hercules CA). Samples were run in triplicate (10μL EvaGreen supermix 2X, 1μL 10μM f primer, 1μL 10μM reverse primer, 1μL (1:100 diluted) template DNA, and 7μL H_2_O; reaction was 95°C for 5 min, [95°C for 10 s, 62°C for 20 s] × 40).

### 16S and ITS2 sequencing

We used a two-step PCR to prepare amplicon libraries as described previously (Wu *et al.*, 2015). For the first step, primer sets used for amplification of the ITS2 and 16S genes were gITS7F/ITS4R (fungal ITS2) (White *et al.*, 1990), and 515F/808R (bacterial and archaeal 16S v4 region) (Supplementary Table 2). Procedures differed from Wu et al. (2015) in amplification cycles (10 cycles in the first step and 20 cycles in the second step for 16S; 12 cycles in the first step and 22 cycles in the second step for ITS). For the second step, phasing primers were used to increase base diversity in sample library sequences. Sample libraries were sequenced on a MiSeq system (Illumina, San Diego, CA, USA) (2x250bp paired ends) at the Institute for Environmental Genomics, University of Oklahoma.

### 16S and ITS sequence data analysis

For processing and analyzing the 16S data, we drew on methods from (Pepe-Ranney *et al.*, 2015; McMurdie and Holmes, 2013). We used Paired End reAd mergeR (PEAR) (Zhang *et al.*, 2014) to merge reads, screed databases (Nolley and Brown, 2015) to demultiplex sequences, cutadapt (Martin, 2011) to remove primers, USEARCH (Edgar, 2013) to filter reads and for OTU clustering (97% ID), mothur (Schloss *et al.*, 2009) for alignment-based quality control, and QIIME (v1.8) (Caporaso *et al.*, 2010) to assign taxonomy, using the green genes 97% ID OTU taxonomy database (details in Supplementary Note 2). For the ITS2 data, we processed them as for the 16S data, drawing on methods from (Bálint *et al.*, 2014; Glassman *et al.*, 2015), with the addition of using ITSx (Bengtsson-Palme *et al.*, 2013) to extract only the fungal ITS2 regions of the reads, and using the UNITE reference database (Kõljalg *et al.*, 2013) at 97% ID to assign taxonomy (details in Supplementary Note 2). We assigned AMF status to any taxa within the *Glomeromycota* phylum, and EMF status to taxa identified as EMF by Glassman et al. (2015). These assignments were largely consistent with the results from using the FUNGuild database at the “highly probable” or “probable” confidence rankings (Nguyen *et al.*, 2016).

### Community structuring processes

To determine the dominant processes structuring communities in the minerals and rhizosphere soils, we used the approach described by Stegen *et al*. (2013). In this framework, ecological processes are classified into the following categories: (i) homogeneous selection (abiotic or biotic pressures select for the same types of characteristics across communities), (ii) variable selection (abiotic or biotic pressures select for different types of characteristics across communities), (iii) homogenizing dispersal (individuals can move between communities easily), (iv) dispersal limitation (individuals can not move between communities easily), and (v) undominated (population fluctuations are essentially due to weak selection, weak dispersal, and/or random chance events) (Stegen *et al.*, 2013; 2015). Briefly, for each pair of communities (samples), variable or homogeneous selection was first inferred in pairings where the phylogenetic dissimilarity (*β* Mean Nearest Taxon Distance, *β*MNTD) was significantly higher (*β*NTI>2, *β* Nearest Taxon Index) or lower (*β*NTI<-2) than null expectations, respectively. In the cases where |*β*NTI|<2, “dispersal limitation” or “homogenizing dispersal” was inferred in pairings where taxonomic dissimilarity was significantly higher (RC_Bray_>0.95, modified Raup-Crick metric based on Bray-Curtis dissimilarity (Chase *et al.*, 2011)) or lower (RC_Bray_<-0.95) than null expectation, respectively. In the remaining cases (|*β*NTI|<2 and |RC_Bray_|<0.95), other stochastic processes were considered to be the governing processes (“undominated”). We used the same null model algorithms as Stegen *et al*. (2013, 2015), but improved the estimation of OTU relative abundances in the metacommunity in the null model as follows: the relative abundance of each OTU in each sample was weighted by estimated biomass amount (according to DNA concentrations) in this type of material in the whole microcosm, to calculate relative abundance of each OTU in the metacommunity. The null model analyses were done within each sampling timepoint, separately. The relative importance of a process was measured as the percentage of comparisons dominated by each process, in all comparisons among communities within soil samples or between mineral and soil samples, for each mineral type. We report the comparisons of each mineral type to soil samples, since we are most interested in the processes that determined how minerals and soil communities were interacting.

In addition to the *β*NTI and the RC_Bray_, we calculated the nearest taxon index (NTI) and net relatedness index (NRI) individually for each sample (Webb *et al.*, 2002), with 1000 randomizations, using the picante package (Kembel *et al.*, 2010) in R (R Core Team). NRI characterizes the mean phylogenetic distance of taxa in a sample from those in all other samples. NTI characterizes the phylogenetic distance from one taxon to the nearest taxon for each taxon in the sample. NRI is a measure of overall clustering, while NTI is more indicative of terminal clustering (Webb *et al.*, 2002).

### Statistical analyses

To determine significant differences between minerals and soil for DNA extractions and qPCR results, we performed single-factor ANOVAs and Tukey’s HSD in R (R Core Team), log-transforming qPCR copy numbers to maintain assumptions of normality. To characterize differences in community composition between samples, we performed a non-metric multidimensional scaling (NMDS) analysis on Bray distances between samples, with OTU counts transformed to relative abundance, using the vegan package in R (Oksanen *et al.*), with k=3. To determine whether the differences in NMDS plots were significant, we performed a permutational multivariate ANOVA on Bray distances using the vegan package in R (Oksanen *et al.*) (Supplementary Notes 2 and 3). To determine which taxa significantly increased or decreased in relative abundance in the minerals, compared to in the soil, we used DESeq2 (Love *et al.*, 2014). We calculated differential abundances for all OTUs for each mineral *vs*. the soil for each timepoint (see Supplementary Notes 4 and 5 for details on outlier OTUs, which DESeq2 excludes based on a Cook’s distance identification of outliers, and independent filtering).

To evaluate possible relationships between differential abundances in the minerals *vs*. the soils and 16S copy number, we predicted 16S copy number for each OTU using the ribosomal RNA number database (rrnDB-4.4.3) (Stoddard *et al.*, 2015; Lee *et al.*, 2009). Briefly, we assigned taxonomic names to our OTUs using the Ribosomal Database Project (RDP) database, searched the rrnDB to determine if that genus was included in the database, and if it was, recorded the mean 16S copy number known for that genus. We note this is only a rough predictor of copy number, since there is known variation even within a single genus, and we were limited by the taxa included in the database. Thus, results should be interpreted with some caution. To evaluate the relationship between 16S copy number and differential abundance, we built a linear model, using copy number and mineral type with an interaction with phylum as predictive factors for differential abundance *vs*. soil using the *lm* function in the R package “vegan” (Oksanen *et al.*).

Because mineral bags were placed in direct contact with growing roots, we expected that the rhizosphere would be a key source of mineral colonizers. To assess whether the bacteria we identified as successful mineral colonizers are generally successful in the rhizosphere, we drew on a previous experiment, where *Avena fatua* (a close relative of *Avena barbata*) was grown in the same soil as this experiment, and bulk and rhizosphere soils were sampled and analyzed over two growing seasons to determine members of the “dynamic rhizosphere” (Shi *et al.*, 2015). Briefly, we took the sequences from the OTUs that were identified as being enriched in any of the mineral samples, and used USEARCH (Edgar, 2013) to cluster the 16S sequences of the OTUs from the dynamic rhizosphere with these mineral-enriched OTUs, using a 97% identity cutoff.

## Results

### Mineral colonization

We extracted significantly more DNA from whole soil than from all mineral types and the heavy fraction (Figure 2). Of the minerals, we extracted significantly more DNA from ferrihydrite and the heavy fraction, and the least from kaolinite. Any kaolinite samples that had DNA extraction and amplification levels below blank controls were excluded from our analyses. These trends were generally mirrored in 16S copies (bacterial and archaeal) (Figure 2) and ITS copies (fungal) (Figure 2), as determined by qPCR. These trends remained similar when considered on a surface area basis (Supplementary Figure 1).

**Figure 2.**
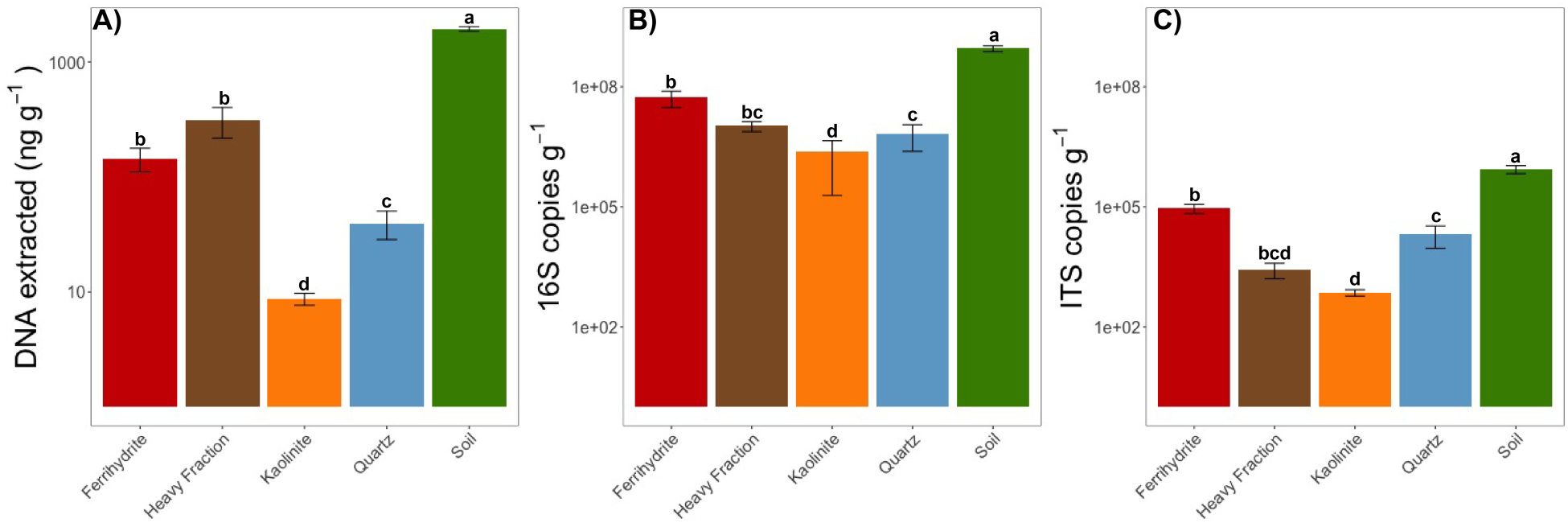
(A) DNA extracted from soils and minerals; (B) 16S copies; (C) ITS copies. Mean values after 2-2.5 months after the start of the experiment, with error bars representing ±SE (n=3 for heavy fraction, n=8-23 for all other minerals). Note log scales. Lowercase letters indicate significant differences (p<0.05, ANOVA, Tukey’s HSD).

### Community comparisons

Community composition differed significantly by mineral type (Figure 3) for both fungi and bacteria/archaea. There was significant effect of mineral type on Bray-Curtis distances between samples for both fungi and bacteria/archaea (Supplementary Figures 2 and 3; permutational multivariate ANOVA, p<0.001). While there were significant changes in community composition over time (p<0.08 for bacteria/archaea and p<0.004 for fungi), these were not dramatic. For the remaining analyses, we present results from the 2 and 2.5 month time-points combined, as they were not significantly different in community composition (p<0.12 for both fungi and bacteria/archaea), had similar levels of diversity (Supplementary Figure 4), and had more DNA extracted than the 1 month time-point (Supplementary Figures 5 and 6).

**Figure 3.**
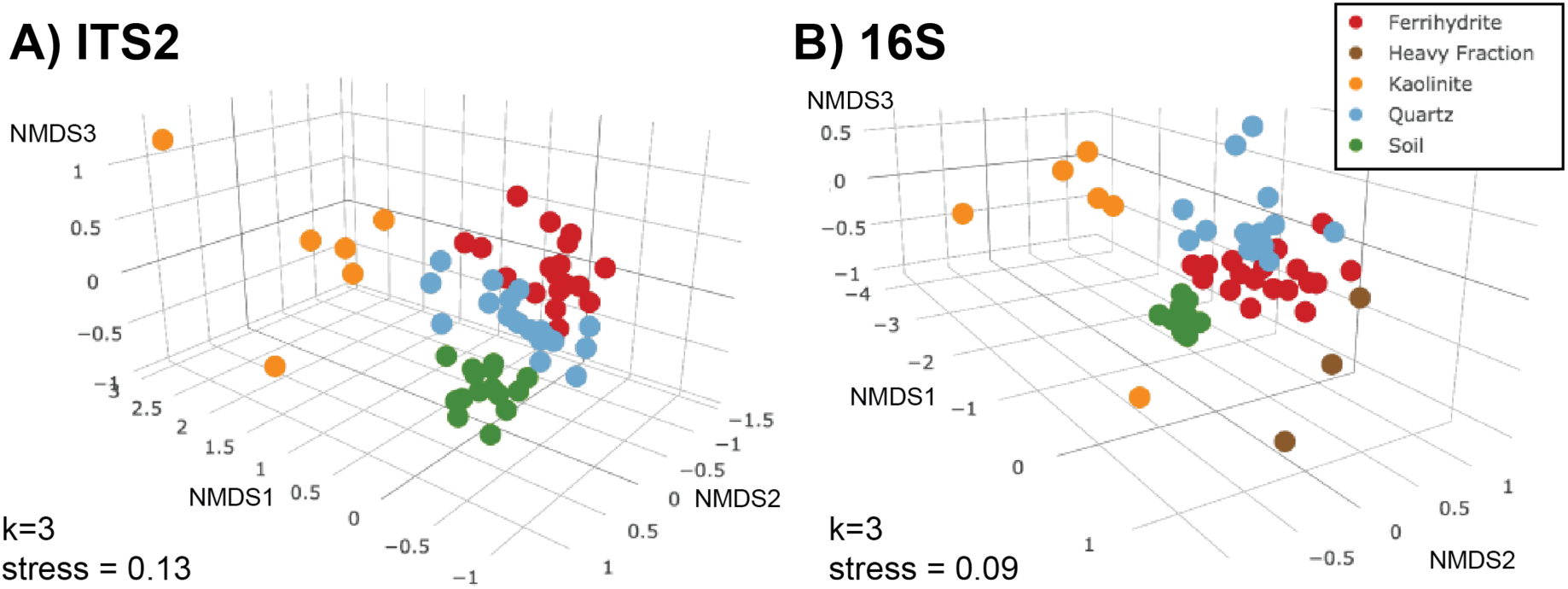
Three-dimensional NMDS plots of Bray distances for A) fungal ITS2 (k=3, stress=0.13) and B) bacterial/archaeal 16S (k=3, stress=0.09) communities, along with bulk soil samples.

### Community composition

At the bacterial phylum level, all samples were dominated by *Proteobacteria*, *Actinobacteria*, *Bacteroidetes*, *Firmicutes,* and *Acidobacteria* (Figure 4). The mineral microbial communities were significantly lower in relative abundance of *Acidobacteria, Planctomycetes,* and *Gemmatimonadetes* than was the soil community.

**Figure 4.**
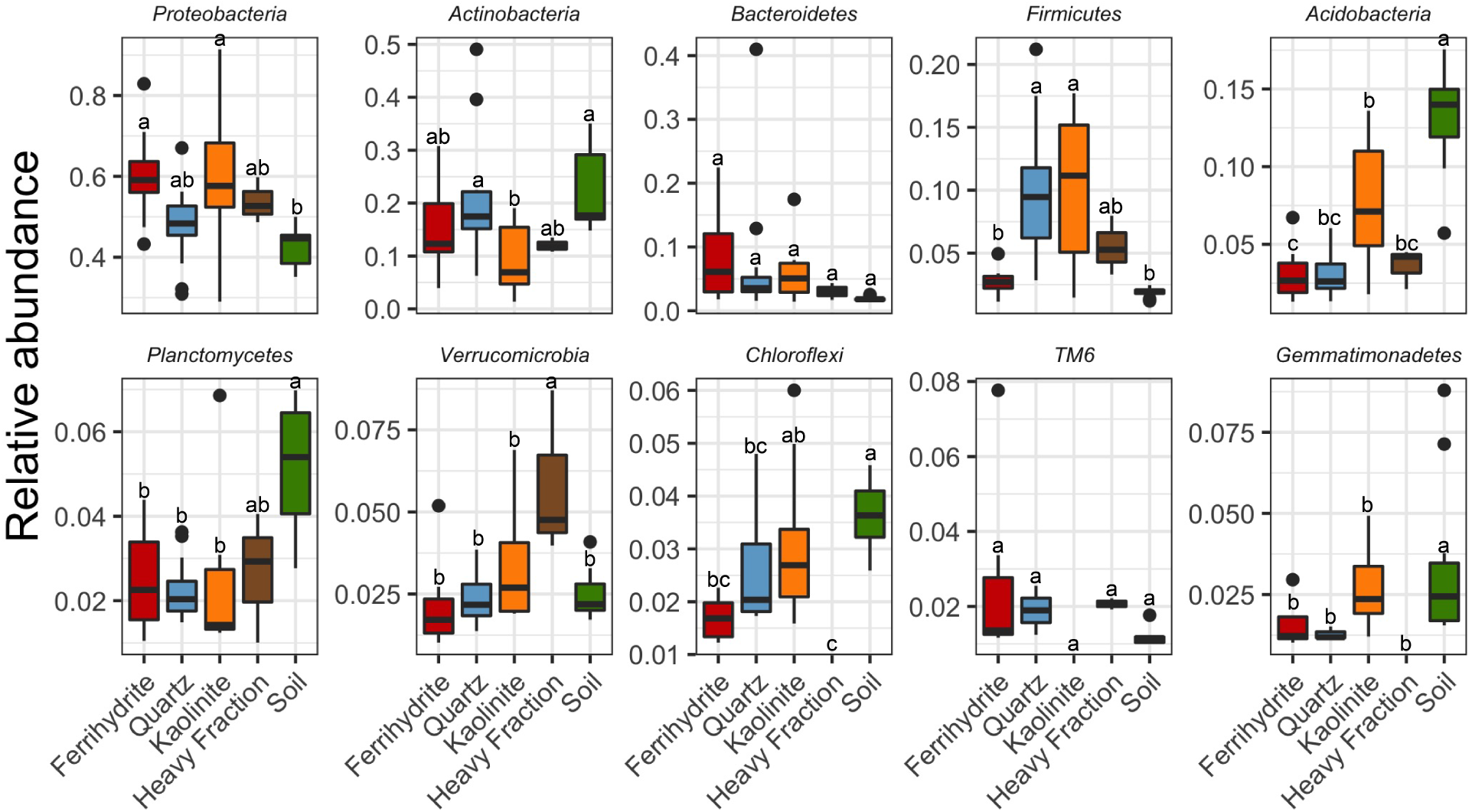
Relative abundance of top 10 bacterial phyla in different mineral types. For heavy fraction, n=3; for all others n=11-14. Letters indicate significant differences within a phylum (p<0.05, ANOVA, Tukey’s HSD).

Fungal communities were dominated by *Ascomycetes* and *Basidiomycetes*, although large fractions (up to 40% in kaolinite minerals) were not identifiable using the UNITE database even at the phylum level. At a finer taxonomic level, the most abundant orders (for those taxa identifiable to order) were *Sordariales*, *Eurotiales*, and *Hypocreales*. The orders *Sordariales* and *Eurotiales* had significantly lower relative abundances in the minerals as compared to the soils, and orders *Sebacinales* and *Glomerales* had significantly higher relative abundances in the ferrihydrite minerals than the soils (Figure 5 and Supplementary Figure 7).

**Figure 5.**
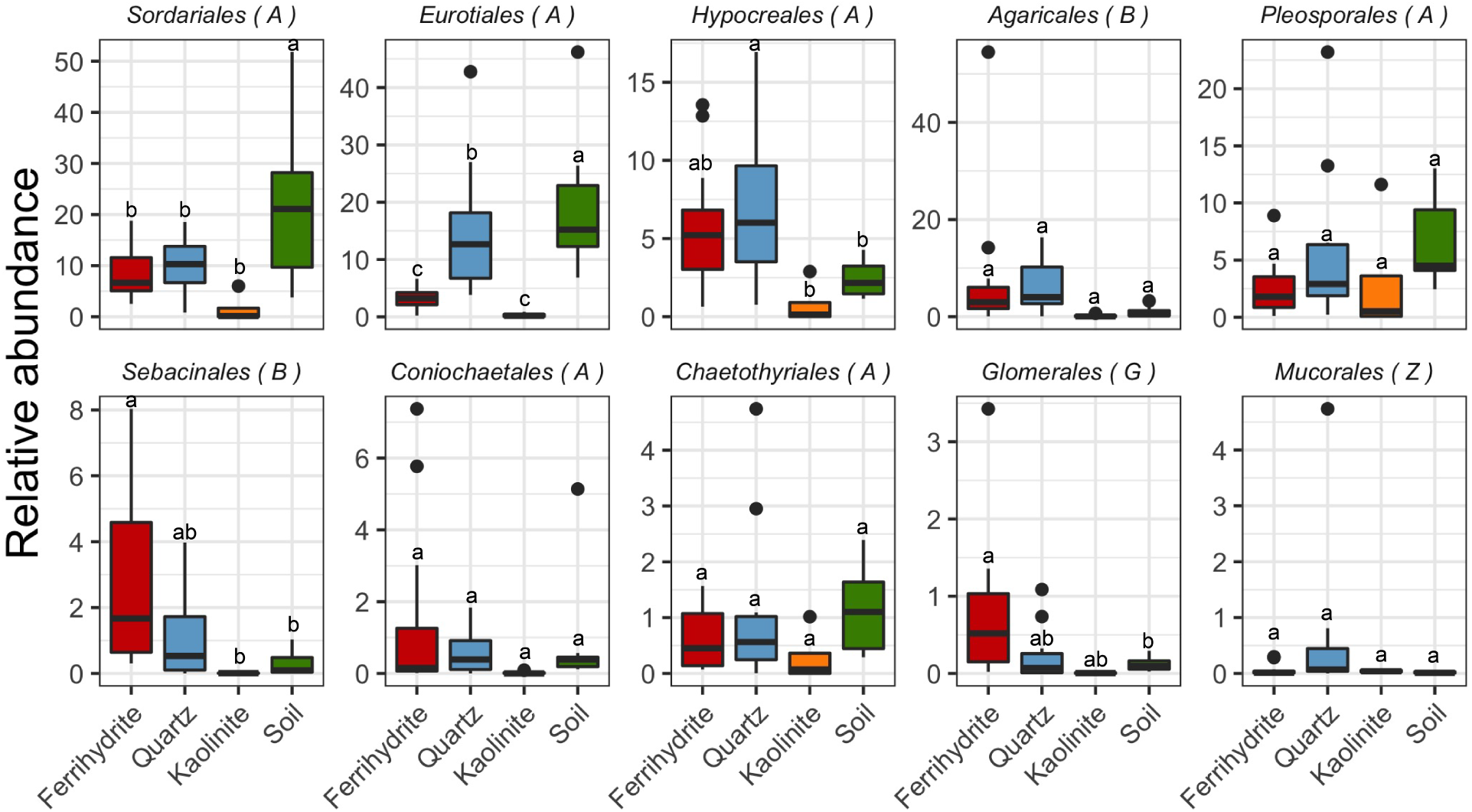
Relative abundance of top 10 fungal orders in different mineral types. Phylum is indicated in brackets: “A” = *Ascomycota*, “B” = *Basidiomycota*, “G” = *Glomeromycota,* “Z” = *Zygomycota*. For kaolinite, n=4; for all others n=10-14. Letters indicate significant differences within an order (p<0.05, ANOVA, Tukey’s HSD).

### Differential abundance

The relative abundance of mineral-associated microbial communities differed from that of the soil community. In the bacteria/archaea, 39% of all OTUs were significantly enriched in at least one mineral as compared to the soil, while 27% were significantly depleted. 75% of all OTUs were neither significantly enriched, nor significantly depleted in any mineral as compared to the soil. *Acidobacteria* tended to be depleted in the ferrihydrite and the quartz minerals, while *Firmicutes* and *Bacteroidetes* tended to be enriched (Supplementary Figures 8 and 9). *Actinobacteria*, *Proteobacteria*, and *Verrucomicrobia* all showed both positive and negative responses. In the fungi, 9% of all OTUs were significantly enriched in at least one mineral as compared to the soil, while 14% were significantly depleted in at least one mineral. 92% of all OTUs were neither significantly enriched, nor significantly depleted in any mineral as compared to the soil. *Glomeromycota*, the arbuscular mycorrhizal fungi, were consistently enriched in relative abundance in the ferrihydrite minerals, but not the quartz or kaolinite (Supplementary Figures 10 and 11), while there was a broader range of responses for the *Ascomycota* and *Basidiomycota*.

In order to examine these responses at a finer phylogenetic resolution, we plotted the OTUs for which there was a significant (FDR<0.1) response of 4x or greater (log_2_-fold change = 2), grouped by family (Supplementary Figure 12). Of these taxa, *Burkholderiaceae, Chitinophagaceae, Comamonadaceae*, *Phyllobacteriaceae, Rhizobiaceae*, *Rhodospirillaceae,* and *Streptomycetaceae* were enriched in both the ferrihydrite and quartz minerals, while and *Bacillaceae* were only enriched in the quartz minerals. In the heavy fraction, we also noted the enrichment of candidate family *Chthoniobacteraceae*, which was dominated by the putative bacterial nematode symbiont *Candidatus Xiphinematobacter sp* (Vanderkerckhove *et al*., 2015).

We were not able to taxonomically resolve the fungi as well as the bacteria and archaea. The ITS2 region diverges at too fine of a genetic scale to match ITS2 sequences to phylogenetic levels coarser than species-or genus-level. Thus, a large fraction (>50% in some samples) of responding taxa were not identifiable taxonomically. Of the identified taxa, OTUs that matched *Serendipita* and *Pochonia* both stood out as strong responders in both ferrihydrite and quartz minerals (Supplementary Figure 13). OTUs that best matched *Agaricales* were also enriched in quartz, while OTUs that best matched *Sebacinales* were also enriched in ferrihydrite. *Trichosporon* was identified as being enriched in the kaolinite minerals.

Prior work suggests the potential for a bacterial taxon to grow quickly or be a strong early colonizer correlates with 16S copy number (Goldfarb *et al.*, 2011Nemergut *et al.*, 2016). 16S copy number was significantly correlated with log_2_-fold change in relative abundance in minerals *vs*. soil, controlling for phylum and mineral type (Supplementary Figure 14) (ANOVA, p<0.0001, R^2^_adj_=0.28). There were not significant interactions between mineral type and 16S copy number.

In order to determine whether the bacteria that we identified as successful mineral colonizers were simply the same subset of soil bacteria that were successful in the *Avena sp.* rhizosphere, we compared our results to those of (Shi *et al.*, 2015), which studied the same soil and *Avena fatua* (a close relative of *Avena barbata*). Of the OTUs that were significantly enriched in the mineral samples, only 8% (kaolinite) to 18% (heavy fraction) were also identified as members of the dynamic rhizosphere (Shi *et al.*, 2015) (Supplementary Figures 15 and 16). One notable difference is the *Firmicutes* phylum, which contains important mineral colonizers, but not notable rhizosphere responders. Of the OTUs common to both the minerals and the rhizosphere, most were from the phyla *Proteobacteria* (50%), *Actinobacteria* (16%), or *Bacteroidetes* (13%) (Supplementary Figure 17).

### Community assembly

The dominant process governing community assembly across the soil 16S communities within a given time-point was homogenizing selection (abiotic or biotic pressures select for the same types of characteristics across communities; Figure 6). Homogenizing selection also played a role in the assembly of quartz and ferrihydrite communities as compared to soil communities. Interestingly, dispersal limitation played dominant roles in controlling community assembly on all mineral surfaces (Fig. 6). However, in contrast to other mineral surfaces, variable selection played a role in governing microbial community structure on kaolinite minerals (Fig. 6). Incorporating OTU relative abundance in the metacommunity null model analyses with Bray-Curtis dissimilarity did not substantially change the trends (Supplementary Figure 18). Mean NTI and NRI both indicated significant phylogenetic clustering for all mineral types (Supplementary Figures 19 and 20).

**Figure 6.**
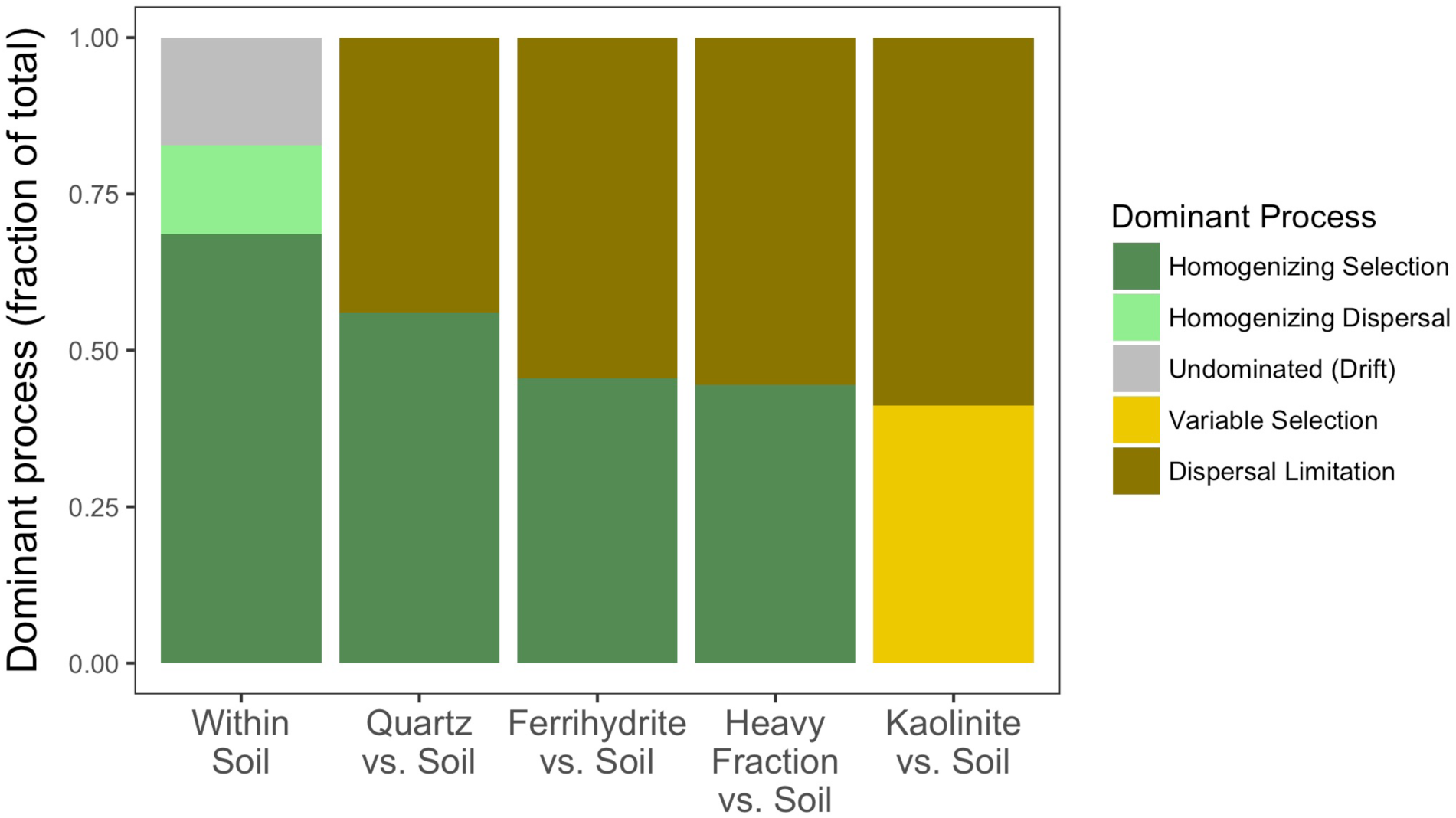
Relative influence of different community assembly processes on spatial turnovers among soil communities and between soil communities and those on different mineral surfaces. The governing processes were determined using RC_Bray_ and *β*NTI (Stegen *et al.*, 2013). Different colors represent different the fraction of community turnovers governed by each process.

## Discussion

### Mineral specificity in microbial community assembly

The mechanisms enabling colonization of soil minerals are likely foundational to the overall soil microbiome composition and partially responsible for the persistence of C entering soil via plant roots. Our results were consistent with previous studies (Wilson *et al.*, 2008; Hutchens *et al.*, 2010; Gleeson *et al.*, 2006; 2005), in that different minerals harbored significantly different bacterial/archaeal and fungal communities (Figure 3), with evidence for selection based on phylogenetic lineage (Supplementary Figures 18 and 19) – *i.e.*, communities have a stronger phylogenetic signal than would be expected by chance. Microbial colonization was likely highest in the ferrihydrite minerals, as we were able to extract significantly more DNA from ferrihydrite than the quartz or kaolinite (Figure 2 and Supplementary Figures 5 and 6). This is also consistent both with other studies, which showed increased microbial biomass on Fe-containing minerals (Wilson *et al.*, 2008), and with our own findings (Neurath *et al*., *in prep*), which showed that ferrihydrite accumulated the most total C. While kaolinite accumulated a comparably high mass of C on a mineral mass basis, when normalized by surface area, it was dramatically lower than that of ferrihydrite (Neurath *et al.*, *in prep*). In addition, the mineral morphology could potentially accentuate the differences in surface accessibility – the platy structure of clay stacks very differently than the granular ferrihydrite or quartz (Neurath *et al.*, *in prep*). The significant differences in the microbial communities that colonize different minerals suggest that mineralogy in natural soils may also be important in determining microbial community structure, with potential implications for biogeochemical cycling and persistence of soil organic matter.

### Possible mechanisms of microbial colonization of fresh minerals

Our null hypothesis was that there would be no meaningful dispersal limitations or selective pressures associated with mineral colonization. The null hypothesis would be consistent with microbes being swept passively onto the minerals with the movements of soil water, and we would have expected that the resulting communities should largely resemble those of the source soil. However, we found that dispersal limitation was an important factor shaping the differences of mineral communities from soil communities, for all mineral types (Figure 6) – that is, in one growing season of an annual grass (2.5 months), a large portion of soil microbes will not be expected reach the minerals by neutral dispersal. Thus, we hypothesize that while this may simply indicate that spatial proximity to minerals may be a key factor in successful colonization, some of the first successful colonizers might be capable of active movement to the minerals. This would require the expression of flagella or other motility factors, such as swarming (Dechesne *et al.*, 2010) and sufficient soil water to support bacterial movement. In addition, water movement by diffusion or advection into and out of the mineral bags could have differed between bags and between bags and the soil. Thus, differences in patterns and processes of water movement may also be responsible for some of the observed differences. Microbes could also be transported to new mineral surfaces by other organisms. For example, we found enrichment of the putative fungal parasite of nematode eggs, *Pochonia* (Supplementary Figure 13) (Kerry and Hirsch, 2011), and the bacterial nematode symbiont *Candidatus Xiphinematobacter sp.* in minerals. While *Candidatus Xiphinematobacter sp.* is thought to be a maternally-transferred obligate symbiont (Lazarova *et al.*, 2016), and so is not likely directly colonizing the minerals, it highlights the potential for movement of soil fauna enabling rapid dispersal of their associated fungi and bacteria.

Once microbes have arrived at fresh mineral surfaces, fast-growing microbes may predominate, by winning the competition for new surface area (Converse *et al.*, 2015). There was a significant positive correlation between enrichment on minerals and predicted 16S copy number (Supplementary Figure 14). 16S copy number has been linked to fast-growth strategies (Goldfarb *et al.*, 2011) and early succession (Nemergut *et al.*, 2016). In addition to fast-growers, organisms that can thrive in environments with low OM may have a colonization advantage, since the minerals are initially low-nutrient environments and high OM may actually inhibit bacterial adhesion to soil particles (Zhao *et al.*, 2014). This low-nutrient, low-OM environment in the mineralosphere could partly explain why there was < 20% overlap in mineral-enriched OTUs with those identified as part of the “dynamic rhizosphere” in a very similar system (Shi *et al.*, 2015) (Supplementary Figures 15-17). While high 16S copy number *Firmicutes* were consistently enriched in the minerals (Supplementary Figures 8 and 9), they were rarely found among the dynamic rhizosphere taxa (Supplementary Figure 17). Thus, some of the characteristics that make a microbe a strong colonizer of new microhabitats in the soil may be different from those that make it a strong responder to roots. For example, while the nutrient environment in the minerals might be expected to reflect that of the rhizosphere in its composition, the total amount of C and other nutrients available were likely dramatically lower in the minerals. One source of nutrients and energy could be the first colonizers of the minerals themselves; possible predatory bacteria such as *Cytophaga* and *Bdellovibrio*, or the possible fungal predator/endosymbiont *Chitinophaga* (Shaffer *et al.*, 2017) were consistently and sometimes dramatically increased in relative abundance in minerals (Supplementary Note 6 and Supplementary Figure 21). Another way to survive in sparse environments could be to access resources from elsewhere via filamentous growth. Significant fungal colonizers of minerals include mycorrhizal symbionts. Unlike saprotrophic fungi, they have a direct plant-derived C source, and so can possibly better “afford” to explore the low-C mineral environments. Supporting this idea, we found that AMF were enriched in ferrihydrite minerals (Supplementary Figures 11 and 22), and *Sebacinales* and *Serendipita vermifera* (possible mycorrhizal fungi) were significantly enriched in ferrihydrite and quartz (Supplementary Figure 12). AMF could potentially act as a sort of shunt of C from the plants to the fresh minerals, paving the way for future mineral colonizers. Mineral-colonizing fungi may directly provide a source of C for bacteria - one of the most mineral-enriched OTUs (log_2_-fold changes of 7.3 and 4.0 in ferrihydrite and quartz, respectively, and representing >1% of the total community in ferrihydrite) was identified as *Chitinophaga sp.* (a possible consumer or endosymbiont of fungi (Shaffer *et al.*, 2017)) and was also part of the dynamic rhizosphere (Supplementary Figure 21). Unlike AMF, we found that predicted saprotrophic fungi (Nguyen *et al.*, 2016) tended to be depleted in ferrihydrite (Supplementary Figure 22), likely because there was little C there on which they could subsist. However, while we might have predicted that fungi would generally be better colonizers of the sparse mineral environments than bacteria/archaea, due to their ability to draw on resources elsewhere through their hyphae, this was not supported by the qPCR data. There were not significant differences in the 16S *vs*. ITS copy number ratios between quartz or ferrihydrite and soils, and the heavy fraction and kaolinite minerals actually had significantly higher 16S *vs.* ITS copy number ratios than were found in soils (Supplementary Figure 23). Furthermore, *Actinobacteria* were not consistently enriched in minerals, although they can often exhibit filamentous growth similar to that of fungi. Thus, filamentous growth alone may not be a reliable predictor of greater colonization success - a robust C source (such as that secured through symbiosis) may also be required.

In addition to dispersal limitation, homogeneous selection was an important factor for all minerals except for kaolinite, which was influenced by variable selection, as compared to the soil communities. While the explanation for homogeneous selection is likely relatively straightforward – certain features of quartz and ferrihydrite resemble those of the soil, and result in similar environments with similar selective pressures – there may be a few possible explanations for the variable selection in kaolinite. These explanations could include various ways in which the kaolinite environment was more different from that of the background soil than were the quartz and ferrihydrite environments, as discussed in the previous section. For example, dramatically higher surface area and smaller particle size (Table 1), may have created an environment substantially different from that of the background soil. A comparatively sparse distribution of resources on kaolinite minerals, given their high surface are, could have resulted in stronger selective pressure for arriving microbes, also contributing to the greater importance of variable selection and competition in these communities (Figure 6). Additionally, the difference in the environments inside vs. the outside of the mineral bags may have been greater for kaolinite than for the other minerals, due to its very small particle size, further differentiating the kaolinite environment.

Our study examines the initial stages of microbial colonization on minerals over a single growing season. Future work following these trends over a longer period of time could address whether dispersal limitation plays a meaningful long-term role in structuring soil mineral communities – do the bacteria that arrive first continue to prevail in the community? This hypothesis may be supported by the observation that the bacterial communities in minerals were more variable than those in the bulk soil (Figure 3 and Supplementary Figures 24 and 25) – suggesting that there is variability in which specific microbes happen to first colonize the fresh minerals. However, only future studies spanning multiple plant growing seasons could determine how long these assemblages might persist, and whether the arrival and establishment of the first sets of microbes could result in exclusion of future potential colonists, or whether the minerals would quickly come to resemble the bulk soil community.

An additional confounding factor is that the soil community represented by ribosomal DNA is an integrated profile of the historical soil microbial community, not just active, or even living, microbes (Blazewicz *et al.*, 2013; Carini *et al.*, 2016). After years in the soil, cycling through disturbances and environmental changes, the apparent (historical) diversity of microbes on the minerals would also be expected to increase, simply as the microbial record of environmental change accumulates. However, bulk soil has more diverse mineralogy than the homogeneous minerals, and, thus, more diverse microenvironments, so some of these differences may persist. For example, we did not seem to see convergence of mineral and soil communities over the 2.5- months of this experiment (Figure 3).

### Conceptual model

While microbial communities associated with the fresh minerals broadly resembled the source soil communities (Figures 2-5), significant phylogenetic differences between mineral and soil communities reveal that community assembly on fresh soil minerals is governed by multiple processes. While we expect passive transport of microbes to fresh mineral surfaces by soil water movements occurs, some microbes are likely actively moving or transported to minerals (Figure 6). Once they encounter the minerals, certain microbes become significantly enriched on the new mineral surfaces (Supplementary Figures 8-11), due to a wide variety of possible biological, geochemical, and physical drivers (Figure 7).

**Figure 7.**
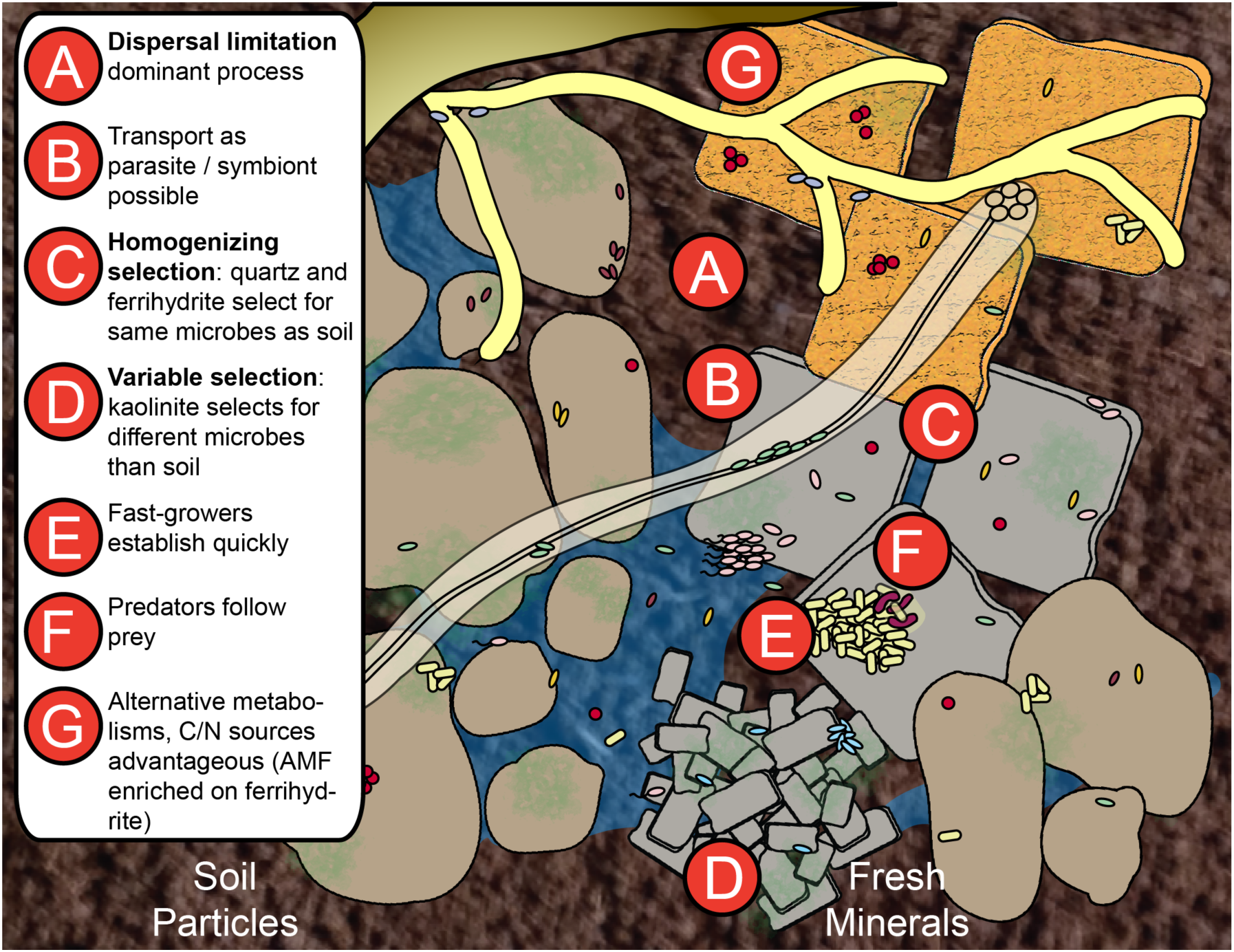
Conceptual diagram of mechanisms by which microbes may colonize fresh soil mineral surfaces. Dispersal limitation (A) was the dominant process over the timescale of this experiment, while some microbes are likely actively moving or being transported to minerals (B). Once microbes encounter the minerals, homogenizing selection structures quartz and ferrihydrite communities (C), while variable selection structures kaolinite communities (D). Fast growth (E), as predicted by predicted 16S copy number, predation (F), and other advantageous factors (G) may result in specific microbes becoming significantly enriched on the new mineral surfaces, due to a wide variety of possible biological, geochemical, and physical drivers.

Mechanisms controlling the colonization of mineral surfaces may be factors in determining the overall composition of the soil microbial community as well as the amounts and composition of biomass and SOC associated with mineral surfaces. To the degree that microbial biomass (and necromass) is an origin of persistent soil organic matter, the dynamics of microbial colonization of soil mineral surfaces are foundational to the stabilization of soil organic matter.

## Acknowledgements

This research is based upon work supported by the U.S. Department of Energy Office of Science, Office of Biological and Environmental Research Genomic Science program under Award Number DE-SC0010570 to UC Berkeley and the University of Oklahoma. Part of this work was performed under the auspices of the U.S. Department of Energy by Lawrence Livermore National Laboratory under contract DE-AC52-07NA27344. The University of California Hopland Research and Extension Center was the source for the soil used here and provided a variety of support services. We thank Katerina Estera, Don Herman, Evan Starr, Shengjing Shi, Erin Nuccio, Anne Kakouridis, and Yonatan Sher for field and lab assistance.

## Supplementary information

Supplementary information is available at XXX. Sequences are deposited in the NCBI short read archive (SRA); the accession number is XXX.

